# Computational Design of Two New Vitamin K Epoxide Reductase Inhibitors

**DOI:** 10.1101/2022.04.06.487400

**Authors:** Chuqiu Cao, Sambid Adhikari, Peishan Huang, Justin B. Siegel

## Abstract

Vitamin K epoxide reductase (VKOR) inhibitors are commonly used to treat atrial fibrillation and ischemic stroke. However, due to the side effects and poor absorbance of existing drugs, there is urgent need for developing better VKOR inhibitor drugs. In this work, computational bioisosteric replacement and chemical intuition were used to design two novel VKOR inhibitors. The new candidates are predicted to have improved ADMET and binding properties compared to existing drugs.

## INTRODUCTION

Atrial fibrillation is the most frequently occurring arrhythmia, characterized by a rapid and irregular heart rate, and triggers a five-fold increase in the risk of ischemic stroke. It is deemed to be the cause of 15% of strokes in the US and greater than 36% for those aged 80 or higher.^1^ The high prevalence of ischemic stroke and atrial fibrillation, especially in the elderly, prompts the pharmaceutical industry to develop drugs with anticoagulation effects, as coagulation has been shown to lead to strokes and atrial fibrillation.^1^ Although originally sold as pesticide, warfarin has since become the most prescribed oral anticoagulation in North America.^2^

By inhibiting the binding of the human vitamin K epoxide reductase (VKOR), the coumarin derived anticoagulant competitively binds to VKOR subunit 1 which is important for activating the vitamin K in the body.^3^ The inhibition irreversibly stops the conversion of vitamin K epoxide to vitamin K1 and the synthesis of Vitamin K dependent clotting factors.^4^ With the depletion of vitamin K reserve, it takes about 2 days for the new coagulation factors to be synthesized.^5^

Several vitamin K antagonists (VKA) have been successful at anticoagulating, such as acenocoumarol, warfarin, phenindione, and chlorophacinone.^6^ Although these drugs have been effective, some pharmacokinetic and pharmacodynamic properties have had adverse effects on patients. They may cause major bleeding in the gastrointestinal tract and mucous membranes if overdosed.^5^ Acenocoumarol even further causes hypotension and peripheral circulatory disorders. ^4^ Warfarin is known as a teratogen and is associated with fetal warfarin syndrome.^5^ Other adverse effects implicated include dorsal midline anomalies, failure of nasal septum development, and underweight can occur in infants.^5^ Hence, new drugs are needed that have less severe side effects.

Studies show that the half-life of R-acenocoumarol (9h) is shorter than warfarin (36-42h), and the half-life of S-acenocoumarol is only 30 min.^7^ Phenindione has a half-life of 5-10 hours^8^ and chlorophacinone has a half-life of about 10 hours.^9^

A few studies have attempted to compare the efficiency of warfarin and acenocoumarol. Pastori et. al. has shown that anticoagulation quality of warfarin is higher than that of acenocoumarol by measuring the mean time in therapeutic range (TiTR) (56.1% ± 19.2% vs. 61.6% ± 19.4%).^10^

According to van der Zee et. al., pharmacogenetic factors also play a role in the dosing of coumarin coagulants. Certain morphisms in VKORC1 and CYP2C9 are linked to lower dosing and higher bleeding, but the clinical effect of genotyping is still under study.^11^ In addition, chlorophacinone is a new longer-acting warfarin derivative and commonly used as a rodenticide. Its initial effect on mats and rice is stronger, but the acute LD50 in *R. norvegicus* is 20.5 mg/kg, which is lower than warfarin. Warfarin-resistant rodents may have no reaction towards chlorophacinone.^12^ Phenindione is now rarely prescribed for its higher probability of severe side effects. It has been associated with liver injuries, such as hypersensitivity and jaundice.^8^ Similarly, acenocoumarol and warfarin are also linked to cholestatic injury.^13^

A previous study proposed a pharmacophore model with three aromatic rings.^14^ The hydrophilic chromene ring that is substituted with a hydroxyl and a carbonyl group is connected to another benzene ring by a one-carbon linkage (Figure 1). With the help of the pharmacophore and existing VKOR inhibitors as a starting point, two novel VKOR inhibitor drugs are designed using the computational and chemical intuition-driven methods in this study. Both candidates exhibited greatly improved predicted oral availability and fewer toxicophore groups based on comparing ADMET (Absorption, Distribution, Metabolism, Excretion and Toxicity) properties to those of the existing drugs. The new candidates also showed improved docking scores suggested they may have improved binding affinities compared to the current VKOR inhibitors warfarin; therefore, they can be served as a possible medium as new treatment for atrial fibrillation.

**Figure 1.**
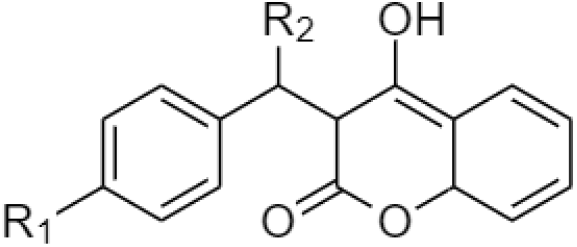
Proposed pharmacophore model, including three aromatic rings, a one-carbon linkage, and a hydroxyl and a carbonyl group that enhances its hydrophilicity.

## METHODS

The crystal structure of human VKOR with inhibitors (PDB ID: 6WV4) bound to the active site was obtained from Protein Data Bank.^15^ All ligand docking simulation is done using OpenEye software suite.^16^ To design a drug candidate based on pharmacophore using the bio-isosteric method, the vBrood application^17^ was used to find replacements of functional groups on the model, acenocoumarol. In addition, SciFinder^18^ is used to confirm the molecule was not previously studied.

Once the ideal candidates were designed, Gaussview and Gaussian 09^19^ were used to build and optimize the drug candidates, respectively. Conformer libraries of known drug molecules and designed drug candidates were generated using Open Eye’s Omega,^20^ after which the conformer libraries of each compound were docked into the active site by using FRED.^21^

Calculatable features that are predictive of ADMET properties, such as molecular weight, hydrogen bond donors and acceptors, and xLogP were found using OpenEye Filter.^22^ PyMol^23^ was used to visualize the protein-ligand interactions as well as finding the measurements between the ligand and the residues.

## RESULTS AND DISCUSSION

### Evaluation of known VKOR inhibitors

The four known drugs acenocoumarol, warfarin, chlorophacinone, and phenindione were built and optimized to evaluate their ADMET properties and docking scores, with the results shown in Table 1. All four drugs satisfy Lipinski’s Rule of Five. Based on the ADMET feature analysis, acenocoumarol is very soluble, phenindione and chlorophacinone are soluble, and warfarin is moderately soluble since acenocoumarol ranks highest in the number of hydrogen bond donors and acceptors (Table 1). Warfarin and chlorophacinone have 3 hydrogen bond donors and acceptors, the second highest. The lowest number of H-bond donors or acceptors explains the moderate solubility of phenindione. The XLogP value of chlorophacinone is 4.64, thus, the compound will need to be administered transdermally, while acenocoumarol, warfarin, and phenindione can be administered orally due to their low XlogP value. The number of hydrogen bond acceptors and donors is the highest in acenocoumarol. It is also the only drug that includes a toxicophore, nitro group. As this toxicophore can be modified for designing new drugs, acenocoumarol would be a good starting point. The ADMET properties show that the drugs have relatively good pharmacokinetics.

**Table 1.** Docking scores and ADMET properties of known inhibitors Acenocoumarol (A), Warfarin (W), Chlorophacinone (C), and Phenindione (P).

|  | Docking Score | Molecular Weight (g/mol) | H-bond Acceptor | H-bond Donor | XLog P |
| --- | --- | --- | --- | --- | --- |
| A | -16.87 | 352.32 | 5 | 0 | 3.04 |
| W | -15.55 | 307.32 | 3 | 0 | 3.12 |
| C | -15.07 | 374.82 | 3 | 0 | 4.64 |
| P | -11.67 | 222.24 | 0 | 2 | 2.66 |

To select one of these VKOR inhibitors as the starting point of drug design, the four inhibitors are docked into the human VKOR active site and the docking scores are listed in Table 1. Phenindione was eliminated as it had the highest value, indicating poorest binding affinity. Warfarin and chlorophacinone didn’t score as negative as acenocoumarol. If improvements can be made on acenocoumarol, the overall score of the known drugs can be further lowered. None of the rules of Lipinski’s Rule of Five is violated, so they are expected to have good absorption and permeation. Taking ADMET properties and docking scores into consideration, acenocoumarol was chosen for drug design.

The interactions between acenocoumarol and the VKOR active site residues were visualized in Pymol, shown in Figure 2. The main interactions observed are the hydrogen bonds between the hydroxyl with ASN222 (2.6Å) and carbonyl with SER165 (3.5Å). The compound is normally charged in neutral pH. The carbonyl on the pyran ring interact with TYR281 via hydrogen bonding (3.1Å). The pi-pi stacking also exists between TYR281 and the two aromatic rings. Based on the docking results the interactions can be exploited to design two new drug candidates.

**Figure 2.**
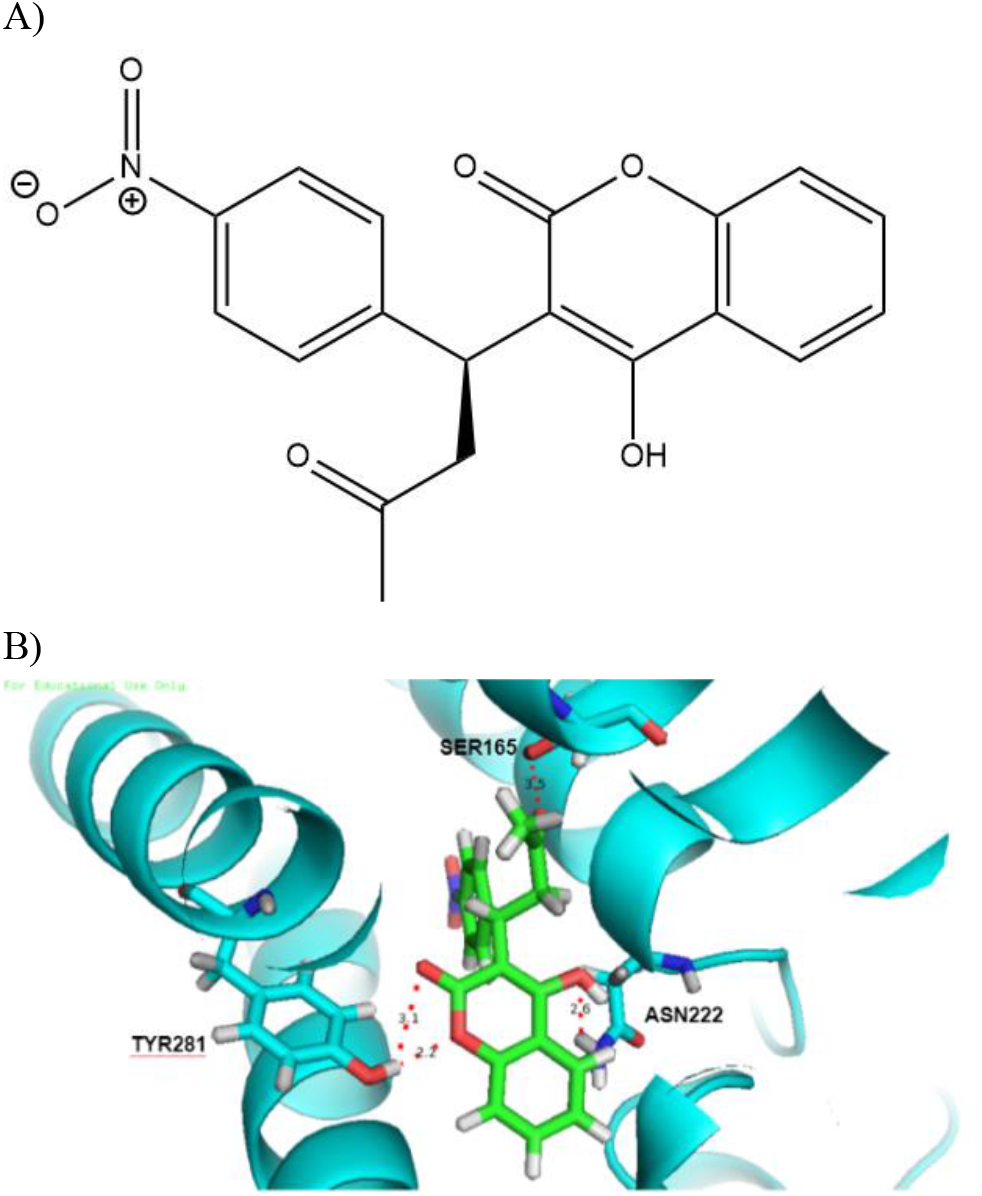
A) 2D structure of acenocoumarol B) Acenocoumarol (green) interactions with SER165, ASN222, and TYR281 (blue).

### Drug candidate 1 designed by computational studies

The first drug candidate was designed computationally by loading acenocoumarol into vBrood. Several candidates that replaced the toxicophore in acenocoumarol were proposed. The conformer library of the candidates that contained the replacement of bio-isosteres were generated and docked into the active site. The conformer with the most negative score −19.92 was named Candidate 1, which was 18.08% higher than acenocoumarol. This molecule included a ketone, a pyrrole that replaced the nitro, and benzene that connected to chromene via a one-carbon linkage. Research has also suggested that a hydrophobic group with hydrogen bond donor at the para position to the carbon linkage of benzene could be key to enhance binding affinity^14^ and potentially explains why the docking score improves.

While most of the interactions of Candidate 1 remain similar with acenocoumarol, its more negative docking score can be explained by a new pi-pi interaction between the pyrrole ring and PHE 229 (5.0Å) (Figure 3B). Additionally, the hydrogen bond between carbonyl and SER165 is 0.4 Å shorter and thereby stronger, compared to that of acenocoumarol. Overall, the new interaction and stronger hydrogen bond contribute to the better binding affinity of Candidate 1.

**Figure 3.**
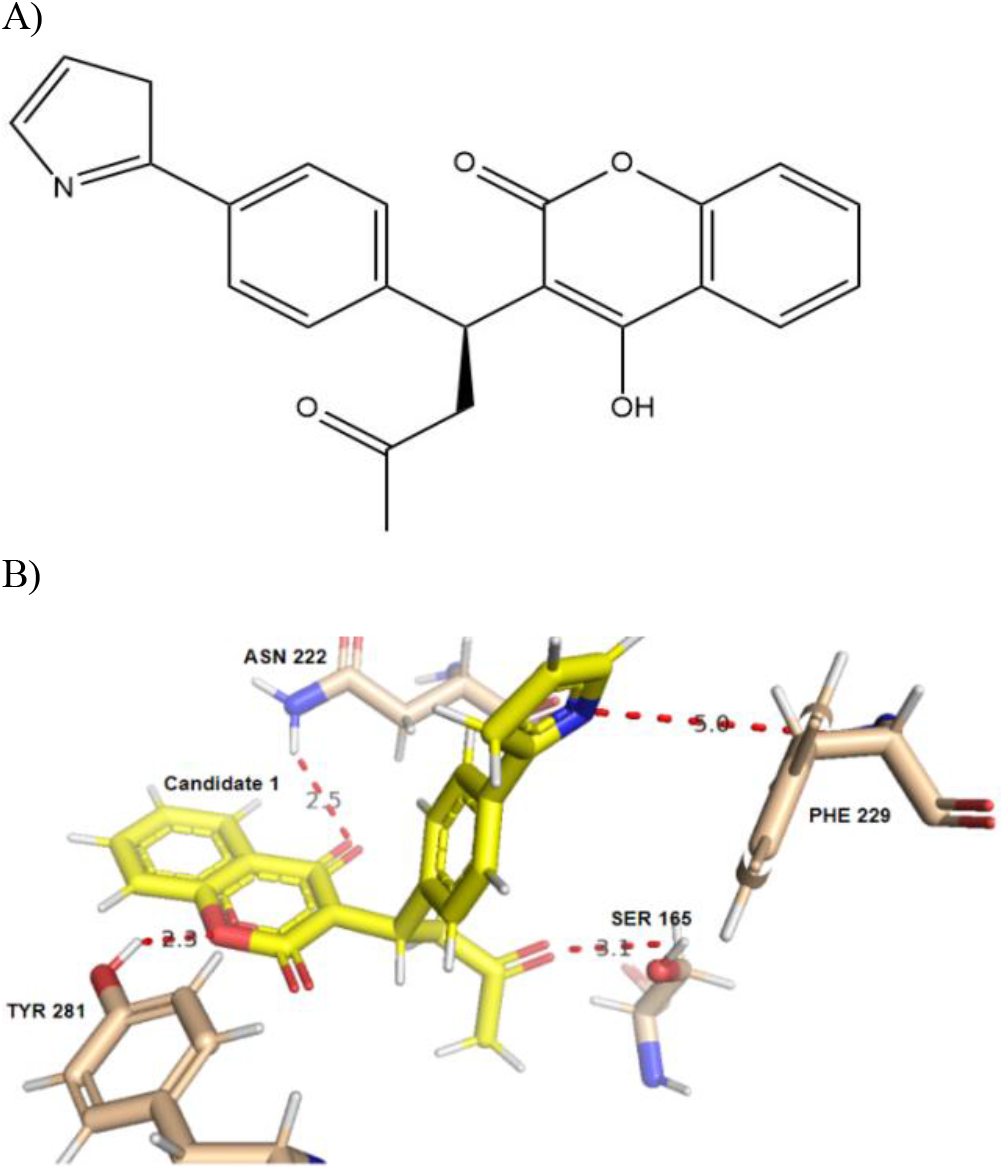
A) 2D structure of drug Candidate 1 B) Candidate 1 (yellow) interactions with SER165, ASN222, TYR281, and PHE229 (wheat).

OpenEye Filter test was carried out to investigate the predicted ADMET properties of Candidate 1. The molecular weight of Candidate 1 is well below 500 g/mol, and its number of hydrogen bond acceptors and donors is acceptable. Its XLogP value is 3.04, which is below 5. Therefore, Candidate 1 is a promising VKOR inhibitor compared to acenocoumarol.

### Drug candidate 2 designed by using chemical intuition

Candidate 1 was subsequently evaluated for further improvements through modifications based on the model in figure 1 and biophysical assessment of potential additional favorable interactions at the protein-ligand interactions at the protein-ligand interface. By incorporating a cyclopropane to the acetate group in Candidate 1, several favorable interactions are predicted to be made with the hydrophobic residues in the active site.

The docking score of Candidate 2 is −20.26, which is 20.09% higher than acenocoumarol. Although the majority of interactions remain the same, Candidate 2 maintained the same interactions as previous control drug (Figure 4B). In addition, Candidate 2 created new hydrophobic interactions show in Figure 4C. The interactions were between the cyclopropane and LEU169 at 3.8 Å, between the benzene ring and PHE197 at 4.1 Å, and ILE265 at 3.8 Å. The hydrogen bond between the carbonyl and SER165, however, was lost, possibly due to the bulky cyclopropane.

**Figure 4.**
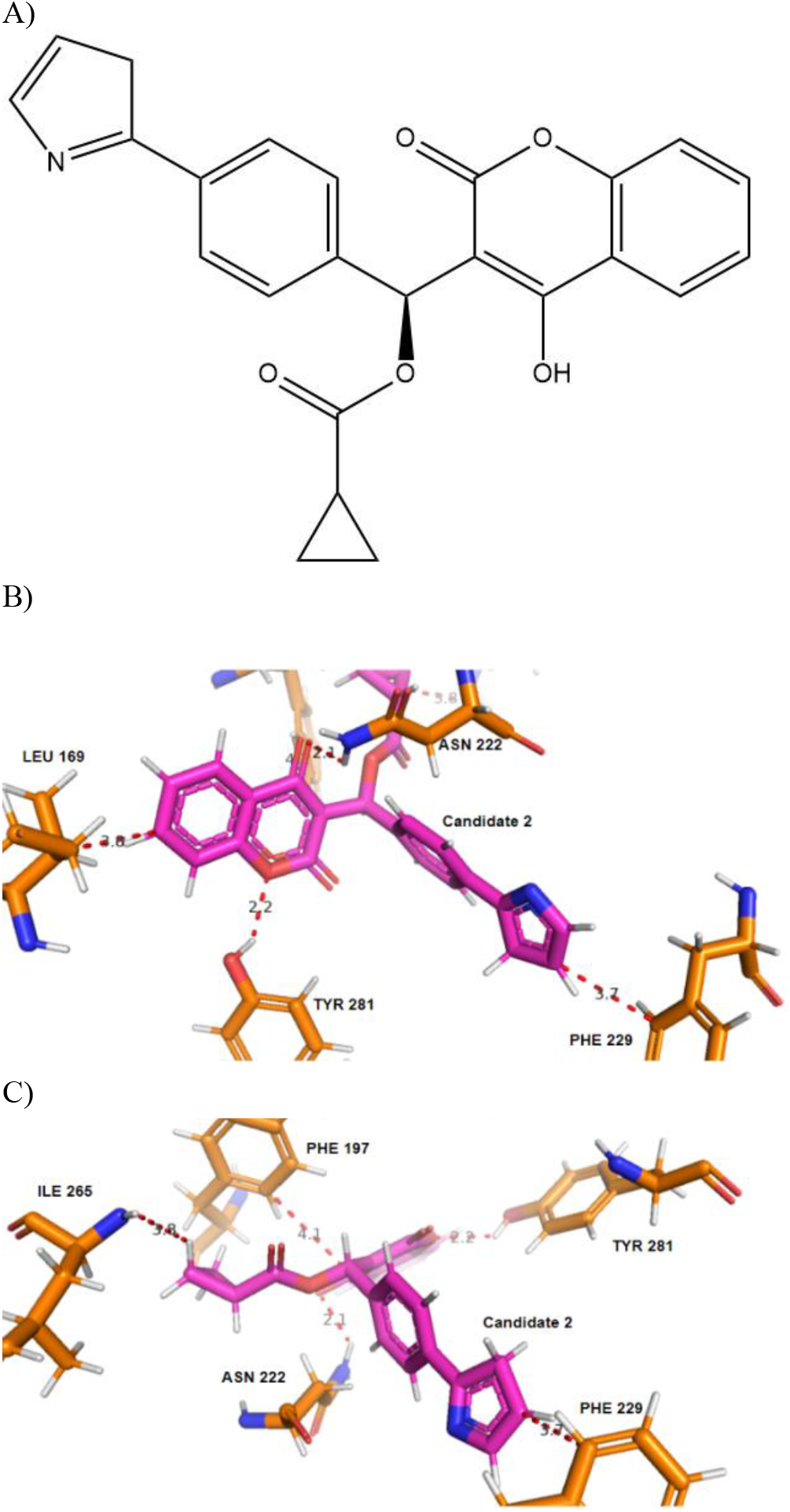
A) 2D structure of drug Candidate 2 B) Candidate 2 (magenta) interactions with ASN222, TYR281, and PHE229, and LEU169 (orange). C) Candidate 2 (magenta) interactions with PHE197, PHE229, ASN222 and ILE265(orange)

The predicted ADMET properties were shown in Table 2. Its molecular weight and the number of H-bond acceptors and donors is in an acceptable range. The number of chiral centers is one. Although the XLogP value is increased, it’s still in an acceptable range to expect to be able to administer orally.

**Table 2.**
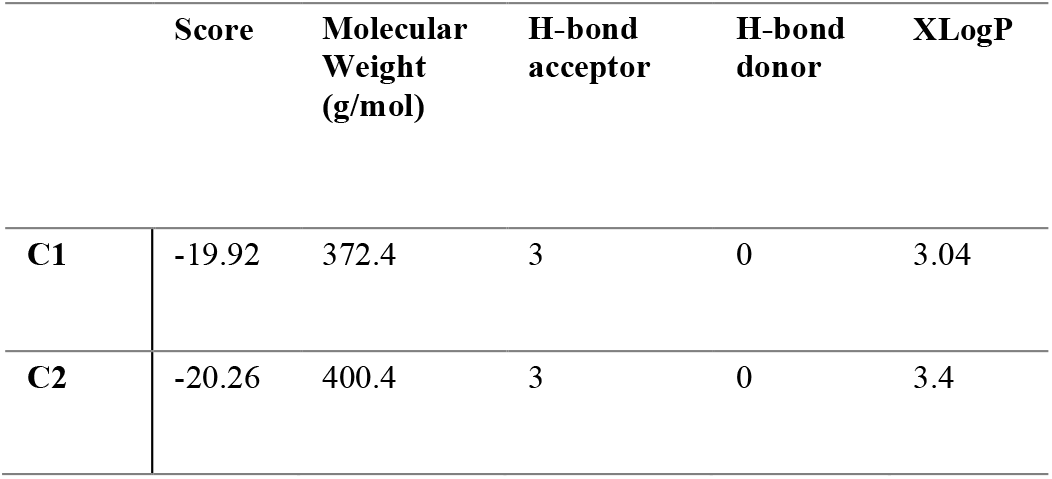
Docking scores and ADMET properties of candidate 1 (C1) and candidate 2 (C2).

## CONCLUSION

With the high rate of ischemic stroke and atrial fibrillation worldwide, anticoagulation drugs are still studied by the pharmaceutical industry. Current VKOR inhibitors on the market have various undesirable adverse effects such as gastrointestinal tract and mucous membranes bleeding. The presence of toxicophores also suggests the need for improvement. In this study, two drug candidates are proposed: one generated by computation and one driven by chemical knowledge, both predicted to have greater binding affinity and meet ADMET requirements.

Although the two candidates are predicted to have promising binding affinity and ADMET properties, further studies need to be done to examine their efficacy and optimize the structure, especially increasing the hydrogen bond donors and acceptors as it’ll improve the strength of hydrogen bonding. The next step is to identify suitable animal models and test the two compounds for safety and efficacy profiles.

## Author Contributions

Research was designed by all authors; all experiments were carried out by C.C. The manuscript was written through contributions of all authors. All authors have given approval to the final version of the manuscript.

## ACKNOWLEDGMENT

Research reported in this publication was supported by UC Davis, the National Science Foundation Award Numbers 1827246, 1805510, 1627539, the National Institute of Environmental Health Sciences of the National Institutes of Health (NIH) under Award Number P42ES004699, UC Davis, NIH Award Number R01 GM 076324-11 and the Rosetta Commons. The content is solely the responsibility of the authors and does not necessarily represent the views of the National Institutes of Health or National Science Foundation. This study was derived from a course based undergraduate research study conducted in Chemistry 130B at UC Davis.

